# ZomB is essential for chemotaxis of *Vibrio alginolyticus* by the rotational direction control of the polar flagellar motor

**DOI:** 10.1101/2021.07.06.451403

**Authors:** Norihiro Takekawa, Tatsuro Nishikino, Kiyoshiro Hori, Seiji Kojima, Katsumi Imada, Michio Homma

## Abstract

Bacteria exhibit chemotaxis by controlling flagellar rotation to move toward preferred places or away from non-preferred places. The change in rotation is triggered by the binding of the chemotaxis signaling protein CheY to the C-ring in the flagellar motor. Some specific bacteria, including *Vibrio* spp. and *Shewanella* spp. have a single transmembrane protein called ZomB. ZomB is essential for controlling the flagellar rotational direction in *Shewanella putrefaciens* and *Vibrio parahaemolyticus*. In this study, we confirmed that the *zomB* deletion results only in the counterclockwise (CCW) rotation of the motor in *Vibrio alginolyticus* as previously reported in other bacteria. We found that ZomB is not required for the clockwise (CW) rotation-fixing phenotype caused by mutations in *fliG* and *fliM*, and that ZomB is essential for CW rotation induced by overproduction of CheY. Purified ZomB proteins form multimers, indicating that ZomB functions as a complex. ZomB may interact with a protein involved in the flagellar rotation, stator proteins or rotor proteins. We found that ZomB is a new player in chemotaxis and is required for the rotational control in addition to CheY in *Vibrio alginolyticus*.

**Importance:** Bacterial chemotaxis is performed by the control of the flagellar rotation. CheY and ZomB control the rotational direction of the flagellar motor in *Vibrio* spp. and *Shewanella* spp. In this study, we characterized ZomB in *Vibrio alginolyticus*, which is essential for the clockwise rotation of the motor.

## Introduction

Many bacteria have flagella, which are the locomotion apparatus of the helical structure. There is a motor at the base of the flagellum in the cell membrane, which converts the influx of H^+^ or Na^+^ into the rotational force of the motor (1–3). The force is generated by the protein-protein interaction between the rotor and stator. The cytoplasmic part of the rotor, which is called the C-ring, is the most important part of the rotor, which is essential for generating torque. The C-ring is composed of three proteins (FliG, FliM, and FliN).

The stator is composed of two proteins, A subunit (MotA or PomA) and B subunit (MotB or PomB). The A and B subunits form an ion channel with arms that bind to the peptidoglycan layer in the cell wall (4–6). The A subunit is a four-transmembrane (TM) protein and the B subunit is a single-TM protein. Recently, cryo-electron microscopy of the stator structure revealed that MotA forms a pentamer surrounding a central axis made of the MotB dimer (7, 8). The MotA pentamer may rotate around the MotB dimer, and the MotA pentamer and C-ring form mesh-like gears that rotate the flagellar motor. The conformational changes of the stator are thought to be mediated by ions passing through the channel in the stator, and the change is converted to rotational motion via electrostatic interactions between the residues in the C-terminal region of FliG and the cytoplasmic region of the A subunit (9, 10). The A subunit has some conserved charged residues that interact with the rotor, and these specific charged residues are located on the outer side of the MotA pentamer structure, farthest from the cell membrane (11).

Most flagellar motors rotate bidirectionally, either clockwise (CW) or counterclockwise (CCW), which allows the cell to control the direction toward a favorable place or away from an unfavorable place. It has been shown that the diameter of the C-ring changes during CCW and CW rotations, and the rotational direction of the motor is reversed owing to the change in the interaction position with the stator (12, 13). In species such as *Escherichia coli* and *Salmonella enterica* that possess peritrichous flagella, CCW rotation of the motors causes the flagellar filaments to form a bundle to push the cell forward, and CW rotation causes the filaments to unwind and pull and tumble the cell. By adjusting the length of time of CCW and CW rotations of the motor, cells can move in the desired direction through a biased random walk (14, 15). In contrast to the bacteria with peritrichous flagella, several polar flagellating bacteria, such as *Vibrio alginolyticus* and *Shewanella putrefaciens*, are pushed forward when the flagellum rotates CCW and pulled backward when it rotates CW.

The structure and mechanism of rotational direction-control of the flagellar motor are well understood in *E. coli* and *S. enterica*. The rotational direction is controlled by the conformational change of the C-ring, which is also called the switch complex (16). In the C-ring, FliG, FliM and FliN assemble to form an inverted cup-like structure in the cytoplasm (17–19). Among them, FliG is located near the TM and directly interacts with the stator. The C-ring is attached to the MS-ring which is formed by 34 molecules of the two-transmembrane protein FliF in the cell membrane (20, 21). FliM binds to FliG, and FliN binds to FliM to form rings away from the membrane in the C-ring (22–26).

Inversion of the rotational direction of the flagellar motor is directed by signals from receptors that sense environmental changes. The inversion is caused by the chemotaxis signaling protein CheY. CheY is converted into phosphorylated CheY (CheY-P), and its phosphorylation is caused by a kinase, CheA, which is activated by signals by chemosensory receptor signals (27, 28). CheY is essential factor for changing the direction of flagellar rotation, and the rotational direction of the flagellum remains CCW in the CheY-deficient strain. CheY-P binds to a well-conserved motif in the N-terminal region of FliM (FliMN) (23, 29–32). CheY-P further binds to the middle domain of FliM or FliN to control the conformational change of the C-ring (23, 33). As a result, a conformational change occurs at the FliG-stator interaction surface in order to switch the rotational direction of the flagellar motor from CCW to CW (34). The CheY-P binding motif in FliM interacts with not only CheY-P but also MinD-like ATPase FlhG which functions as a regulator of the number of flagella in some bacteria such as *S. putrefaciens* and *V. alginolyticus* (35).

The single-TM protein ZomB, whose molecular weight is ca. 30,000, is essential for flagellar motor switching in some bacterial species, including *S. putrefaciens* and *V. parahaemolyticus* (36). The deletion of *zomB* locks the flagellar rotation in the CCW direction which is similar to the *cheY* deficient mutant. Even though *E. coli* does not have the ZomB ortholog, *E. coli* cells overproduce ZomB to increase the switching frequency of flagellar rotation, suggesting that ZomB affects the flagellar motor in common mechanisms among different bacteria (36). The loss of the N-terminal TM region or the highly conserved C-terminal motif (KKKW) causes ZomB to lose its function (36). Overproduction of ZomB decreases the swimming ability of bacteria placed in soft-agar plates, which is not due to the defects in the growth speed or the swimming speed of the cells, suggesting that ZomB affects chemotaxis (36). The chemotactic system is localized to the cell pole also in the Δ*zomB* mutant cells as in wild-type cells, and even the introduction of the gain-of-function mutant CheY does not lead to the CW directional rotation in the Δ*zomB* mutant (36), indicating that ZomB acts directly or indirectly to induce or sustain the CW state of the C-ring conformation. In *S. putrefaciens*, the Δ*zomB* phenotype was suppressed by the deletion of the C-terminal 14 amino acid residues of FliM (36). The C-terminal region of ZomB has been shown to interact directly with the C-terminal region of HubP, which is a conserved protein in all *Vibrio* spp. and *Photobacterium* spp., and works as a polar landmark anchoring various ParA-like proteins to the cell pole (37). In *V. alginolyticus,* HubP defects increase the polar flagellar number, which is similar to the *flhG* mutant (38).

In this study, we investigated ZomB in *V. alginolyticus* and revealed that the deletion of *zomB* led to a complete loss of chemotaxis in the cells. We showed that ZomB is essential for the rotational direction control of the flagellar motor in *V. alginolyticus*.

## Results

### The *zomB* mutation caused CCW-locked rotation of the polar flagellum in *V. alginolyticus*

We prepared a Δ*zomB* mutant (NMB364) of *V. alginolyticus* VIO5, which has a wild-type polar flagellum. We found that the mutant completely lost the motility in the soft-agar plate (Fig. 1A). When *zomB* was overexpressed in the *V. alginolyticus* VIO5 (wild-type), the motility ring in the soft-agar plate was slightly smaller than that in the wild-type without overexpression. Swimming speed and switching frequency of the flagellar motor were measured using a dark-field microscope (Fig. 1B and 1C). In the Δ*zomB* strain, the flagellar rotation was fixed to CCW, the bacteria swam with the flagellum behind, and the rotational direction remained unchanged (Fig. 1C). The swimming speed was faster than that of the wild-type (Fig. 1B). Expression of *zomB* from a plasmid in the Δ*zomB* strain conferred a change in the direction of swimming, although the switching frequency did not recover to the level of the wild-type (Fig. 1C). It is known that the switching frequency of the motor and the proportion of the CW direction rotation increase when the *cheY* gene is overexpressed in wild-type cells (39). We found that none of the Δ*zomB* mutants conferred a CW rotation direction even when *cheY* was overexpressed (Fig. 1C). These results were similar to those of a previous study on *S. putrefaciens* and *V. parahaemolyticus* (36).

**Fig. 1.**
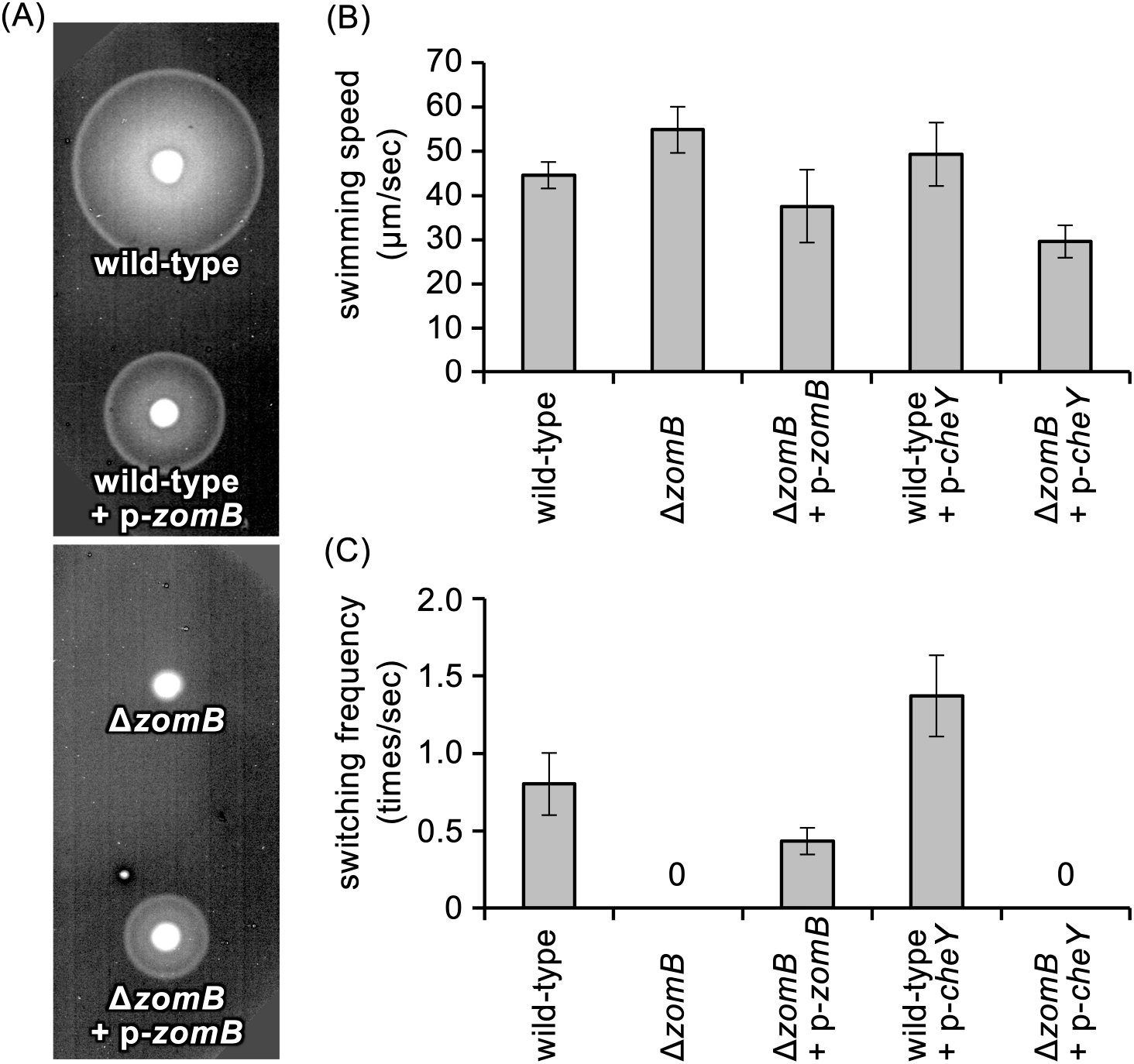
The effect of ZomB on the motility of *Vibrio* cells. (A) The cell motility in soft-agar plate. The overnight cultures of VIO5 cells containing pMMB206 vector (wild-type) or pTN73 (wild-type + p-*zomB*), and NMB364 containing pMMB206 vector (Δ*zomB*) or pTN73 (Δ*zomB* + p-*zomB*) were spotted onto VPG soft-agar plate and incubated at 30 °C for 5 hours. (B and C) The swimming speeds (B) and the frequency of switching of the rotational direction of the motor (C) of VIO5 (wild-type), NMB364 (Δ*zomB*), NMB364 containing pNT73 (Δ*zomB* + p-*zomB*), VIO5 containing pHIDA6 (wild-type + p-*cheY*), and NMB364 containing pHIDA6 (Δ*zomB* p-*cheY*).

### Properties of purified ZomB protein

ZomB is a single-TM protein whose TM region is located at its N-terminus, and the amino acid sequence identity and similarity between ZomB of *V. alginolyticus* and *S. putrefaciens* are approximately 30% and 70%, respectively (Fig. S1). The predicted secondary structure of ZomB is rich in α-helices (Fig. S2A). When the His_6_-tag was fused to the N-terminus or C-terminus of ZomB and expressed in the Δ*zomB* mutant (Fig. S1B), both strains were motile in soft-agar plates, although the diameter of the motility rings was slightly inferior to that of the wild-type cells (Fig. 2B), indicating that the addition of the His_6_ tag had little effect on the function of ZomB. The effects of the deletion of TM region (ZomB_ΔTM_; Fig. S2B) or the C-terminal highly conserved KKKW motif (ZomBΔC; Fig. S2B) on flagellar motility were also examined (Fig. 2A). The ZomB_ΔTM_ strain lost motility in soft-agar plate, as in the case of *S. putrefaciens* (36). In contrast, the ZomB_ΔC_ strain conferred the better motility in soft-agar plate, which is different from the case of *S. putrefaciens* (36).

**Fig. 2.**
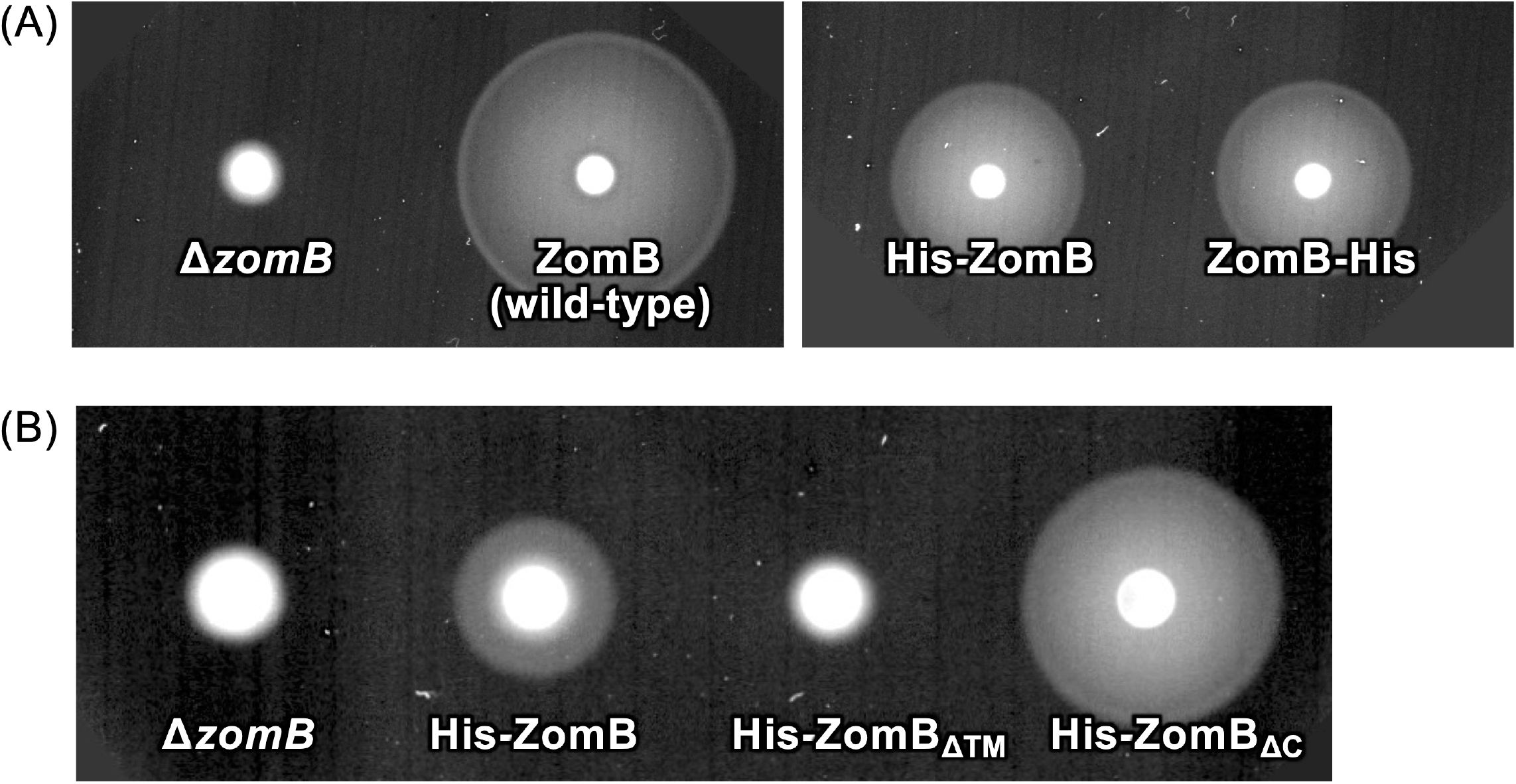
Motility of the cells expressing *zomB* with various mutations. (A and B) The results of the motility assay in the soft-agar plates. The overnight culture of NMB364 containing pMMB206 vector (Δ*zomB*), pNT74 (His-ZomB), pNT77 (ZomB-His), pNT75 (His-ZomB_ΔTM_) or pNT76 (His-ZomB_ΔC_) was spotted onto VPG soft-agar plates and incubated at 30 °C for 7 h (A) and 5 h (B).

The ZomB protein without TM region and with N-terminal His_6_-tag (His-ZomB_C_) was overproduced in *E. coli* and purified by using Ni-affinity chromatography and size-exclusion chromatography. His-ZomB_C_ is expressed as a soluble protein. When the Ni-affinity purified protein followed by removal of the His_6_-tag by thrombin digestion was subjected to the size-exclusion chromatography, it was separated into peaks at approximately 25 kDa, 50 kDa, and 75 kDa, and higher (Fig. 3A). The proteins in each peak fraction were detected as a single band with approximately 30 kDa using sodium dodecyl sulfate-polyacrylamide gel electrophoresis (SDS-PAGE) followed by Coomassie Brilliant Blue (CBB) staining (Fig. 3B), indicating that purified ZomB_C_ proteins formed multimer complexes. We crystallized the ZomB_C_ proteins using the peak fractions (f and g in Fig. 3A), and obtained the two crystal types (Fig. S2A and S2B). These crystals X-ray diffracted at resolutions of up to 3.1 and 2.8 Å (Fig. S2). Both crystals belonged to the space group *P*2_1_ with similar but different unit cell dimensions. The Matthews coefficient (V_M_) value (40) suggested that the crystal asymmetric units contain one molecule with a solvent content of approximately 40 %. We have prepared the Se-Met derivative crystals for phase determination, but no significant anomalous difference was observed in the diffraction data, although the XAFS measurement indicated the incorporation of Se-Met in ZomB_C_. Thus, we could not determine the structure of ZomB_C_.

**Fig. 3.**
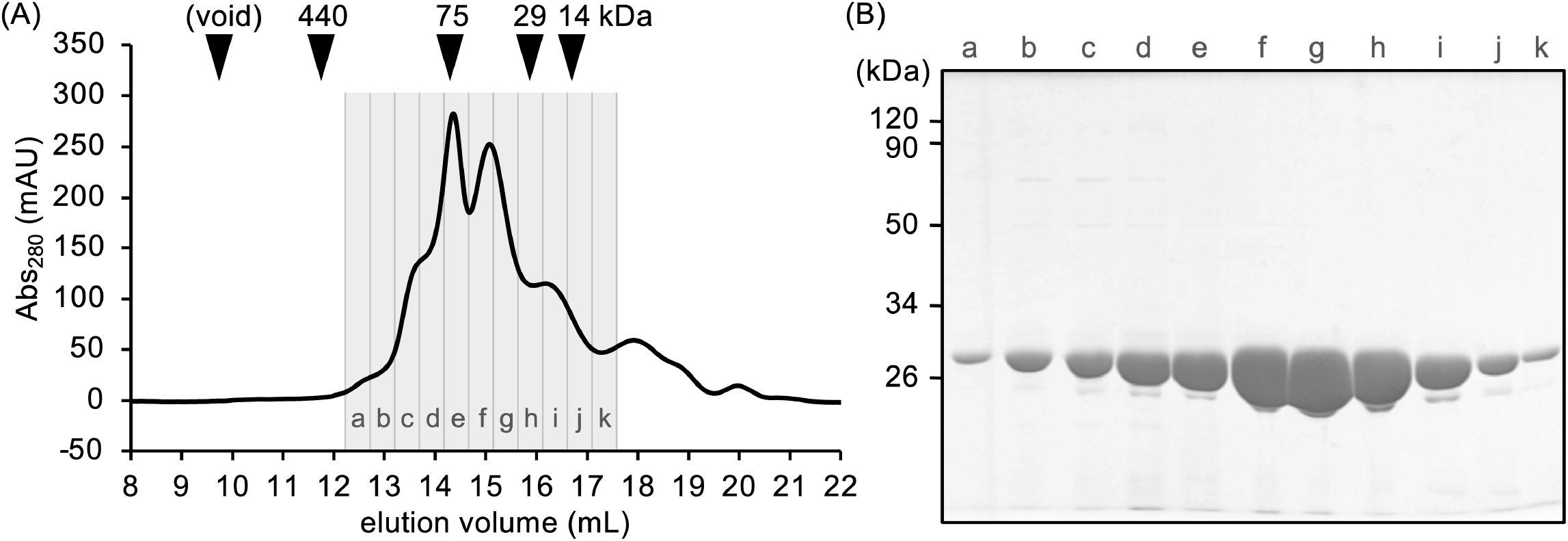
The purification profile of ZomB_C_. (A) Elution profile of purified ZomB_C_ protein by size exclusion chromatography. The vertical and horizontal axes indicate the the absorbance at wave length of 280 nm and the elution volume, respectively. (B) The protein fractions in (A) subjected to SDS-PAGE and stained by CBB.

### Epistasis of the CCW-locked flagellar rotation by Δ*zomB* mutation against the CW-locked flagellar rotation by repellent or mutations in *fliG* and *fliM*

The Δ*zomB* mutation causes the flagellum to rotate only CCW. We investigated the control of the rotational direction in the presence of a repellent, phenol. It is known that addition of phenol extremely increases the switching frequency in the wild-type VIO5 cells (Fig. 4A and 4B) (41). Here, the addition of phenol did almost not affect the motility of Δ*zomB* cells, but a few cells were observed swimming by CW flagellar rotation (Fig. 4A and 4B).

**Fig. 4.**
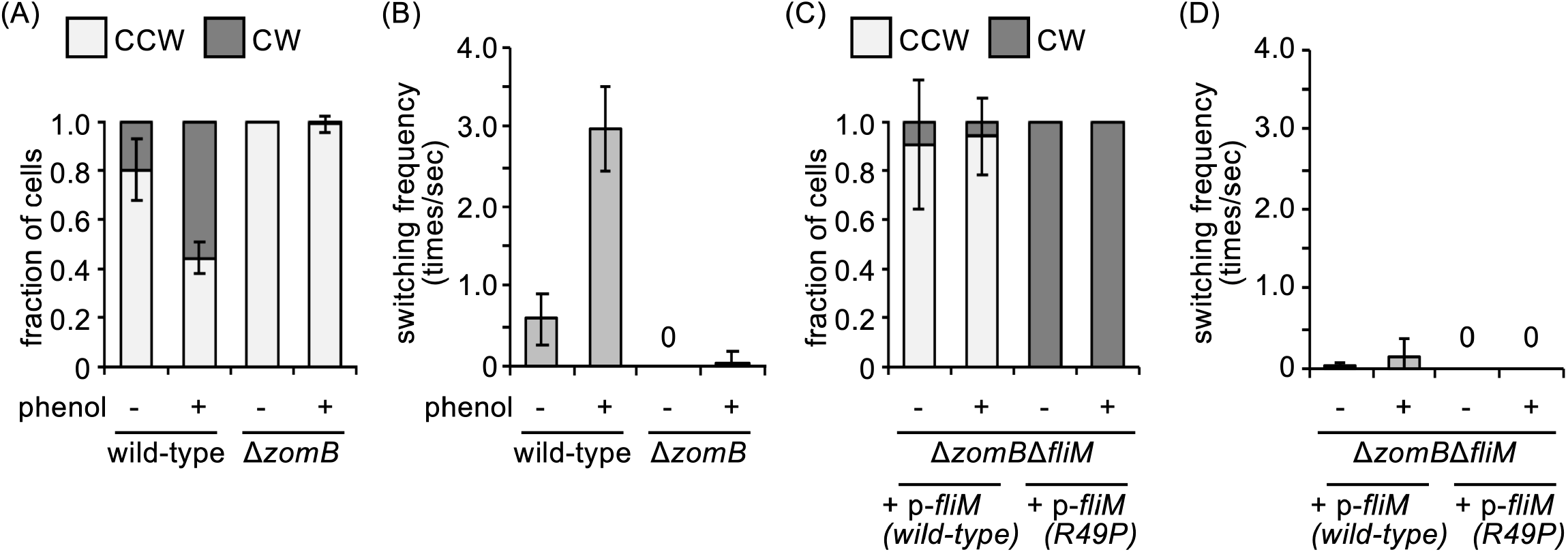
The effect of a repellent (phenol) and *fliM* mutation causing CW rotation to Δ*zomB* mutant. (A) The rates of the rotational direction of the flagellar motors with/without the mutations and phenol. VIO5 (wild-type) and NMB364 (Δ*zomB*) cells with/without phenol were observed using high-intensity dark-field microscopy, and the ratios of CCW rotation (pale gray bar) and CW rotation (dark gray bar) were obtained. (B) The frequency of switching of the rotational direction of the motor. Cells prepared as (A) were observed. (C) The rotational direction rates measured as in (A). NMB365 cells containing pMK1001 (p-*fliM*) with no mutation (wild-type) or R49P mutation were observed. (D) The switching frequency measured as in (B). Cells prepared as (C) were observed.

We also observed the motility of the double mutant of *zomB* and *fliM* (Fig. 4C and 4D). We created a Δ*zomB* Δ*fliM* strain (NMB365) from the wild-type VIO5 strain and introduced a plasmid expressing *fliM* with or without mutation. It is known that R49P substitution in FliM completely locks the rotational direction of the motor in the CW direction (Fig. 4C and 4D) (41, 42). The substitution of FliM-R49P in Δ*zomB* completely locked the flagellar rotation in the CW direction (Fig. 4C and D).

Next, we investigated the motility of the double mutants of *zomB* and *fliG* (Fig. 5). It is known that Q147, G215A, and A282T substitutions in FliG confer CW rotation to the flagellum (43). We created a Δ*zomB* Δ*fliG* strain (NMB366) from the wild-type VIO5 strain and introduced the plasmids expressing *fliG* with or without the mutations (Fig. 5). The CW rotation by G215A mutation was not affected by Δ*zomB*, but the CW rotation by Q147H or A282T mutations changed into the CCW direction by the Δ*zomB* mutation (Fig. 5).

**Fig. 5.**
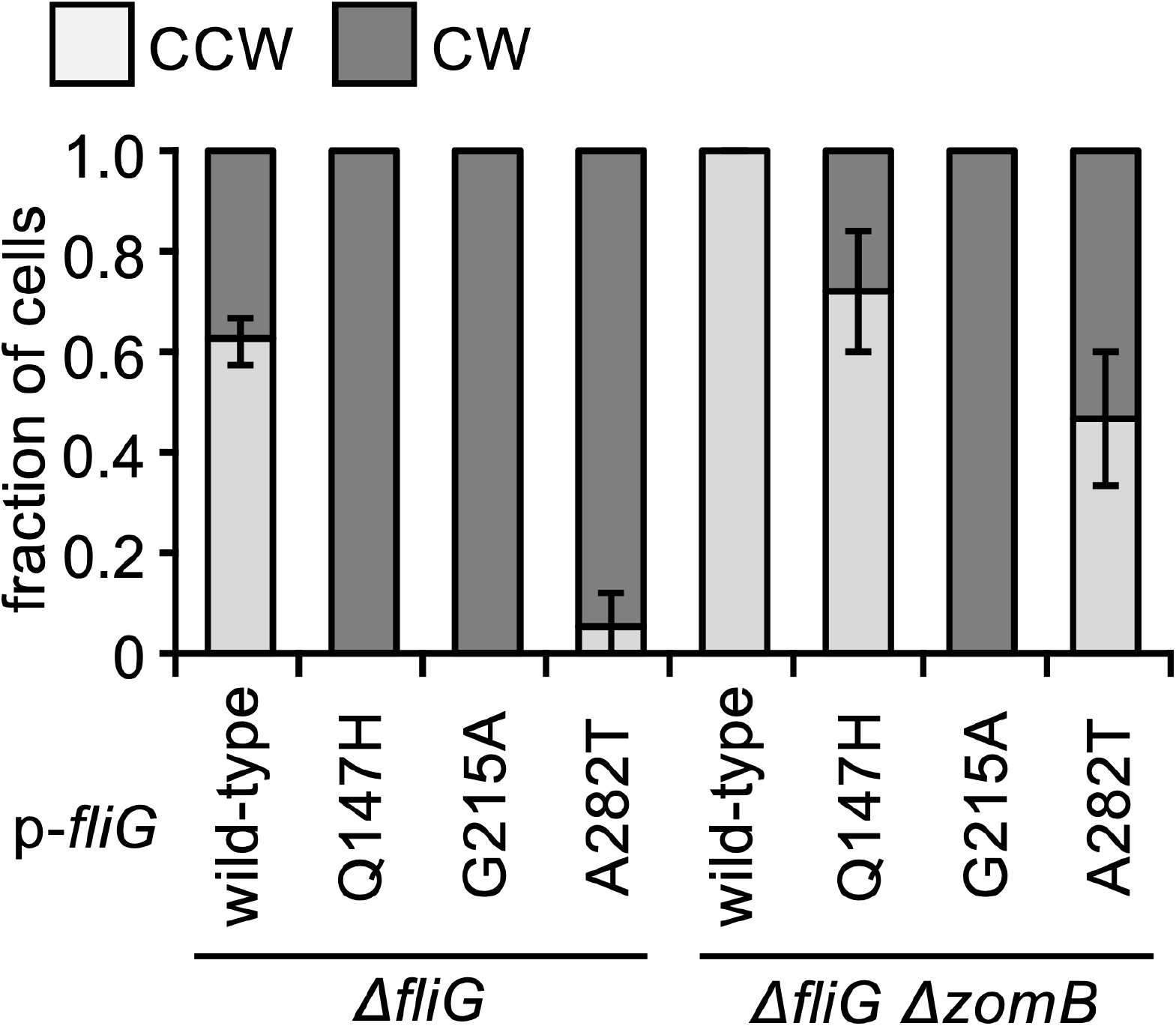
The effect of *fliG* mutations causing CCW or CW rotation in the Δ*zomB* mutant. The rotational direction rates measured as in Fig. 4A. NMB198 (Δ*fliG*) or NMB366 (Δ*fliG* Δ*zomB*) cells containing pNT1 with no mutation (wild-type), FliG-Q147H, G215A or A282T mutations were observed.

### Effect of the C-terminal deletion of FliM in Δ*zomB* mutant

It is known that the CCW-locked phenotype by deletion of *zomB* is suppressed by the deletion of the C-terminal 14 amino acid residues in FliM in *S. putrefaciens* (36). Here we attempted to confirm whether there is a similar effect in *V. alginolyticus*. We produced wild-type or mutant FliM proteins that lacked the C-terminal 10, 20, 30, and 36 amino acids from the plasmids in the Δ*fliM* strain (NMB366). First, we found that overproduction of FliM from the plasmids increased the rate of the CCW rotation as compared to that in the wild-type VIO5, which expresses FliM from chromosomes; approximately 80% in VIO5 but about 90% in the strain overproducing FliM (Fig. 4A and 6A). We also found that overproduction of FliM increased the switching frequency of the motor as compared to that of the wild-type VIO5: approximately 0.7 times per second in VIO5 but about 1.2 times per second in the FliM overproducing strain (Fig. 4B and 6B). The reasons for the effects of overproduction are unknown. However, the overproduced FliM might interact with CheY. Next, we confirmed that all the mutants formed flagella similar to the Δ*fliM* expressing wild-type FliM and were sufficiently motile, as observed under the microscope. The FliM_ΔC10_ and FliM_ΔC20_ mutants were motile in the soft-agar plate. However, the FliM_ΔC30_ and FliM_ΔC36_ mutants were not sufficiently motile (Fig. 6C). The FliM_ΔC20_ and FliM_ΔC30_ producing cells conferred CW-biased flagellar rotation as compared to the cells producing wild-type FliM (Fig. 6A). When the C-terminal deletion FliM proteins were produced in the Δ*zomB* Δ*fliM* strain (NMB365), none of the mutants were motile in soft-agar plate (Fig. 6D), and the flagellum almost lost its switching ability, although the CCW and CW fractions were similar to the case without Δ*zomB* (Fig. 6A and 6B). These results indicated that the C-terminal deletion of FliM did not suppress the CCW-locked phenotype of the deletion of *zomB* in *V. alginolyticus*, which is different from the result of *S. putrefaciens*.

**Fig. 6.**
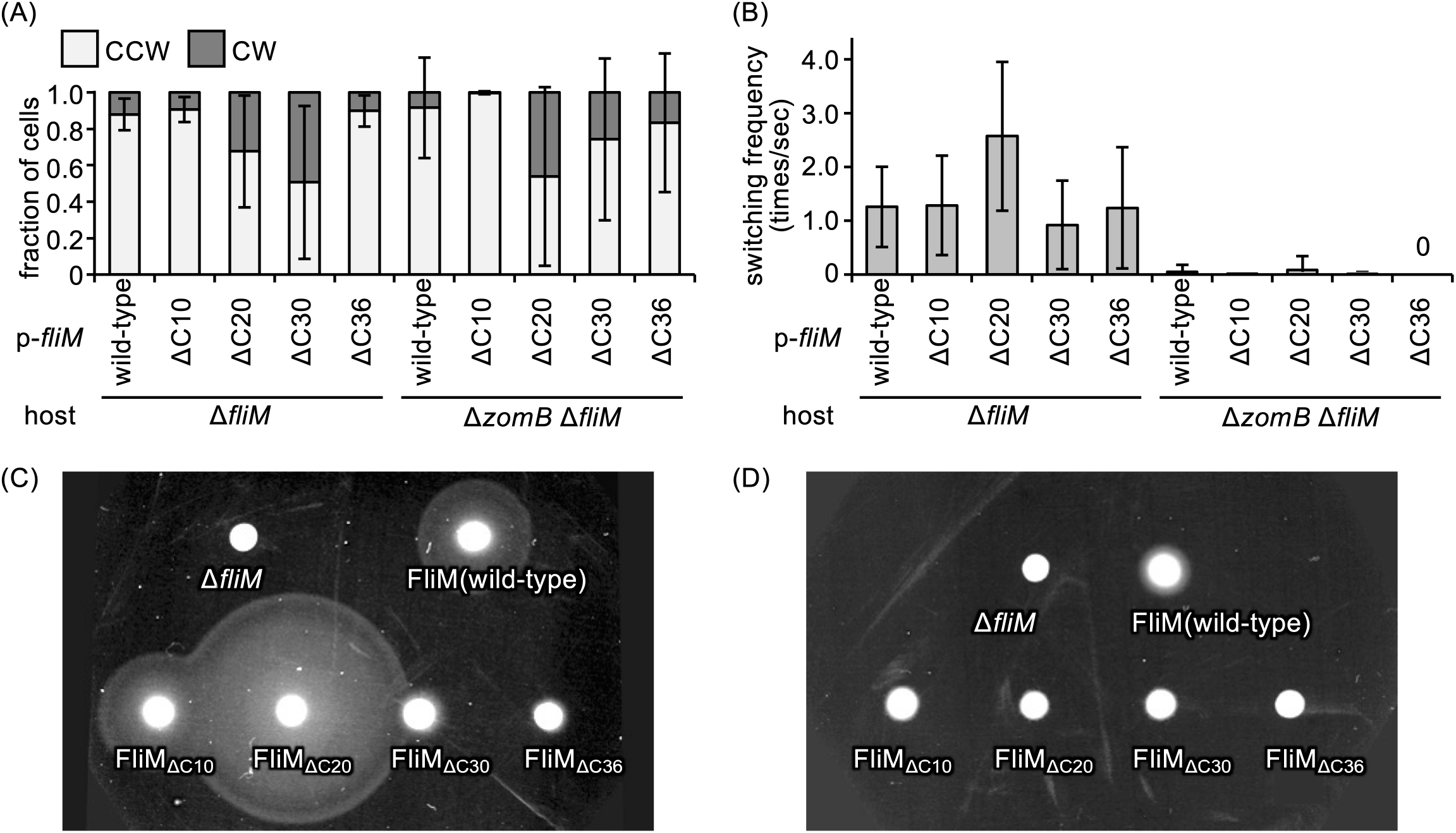
The effect of C-terminal deletion of FliM. (A and B) Motility of the mutants in soft-agar plate. The overnight culture of NMB321 (Δ*fliM*; A) or NMB365 (Δ*fliM* Δ*zomB*; B) containing pBAD33 vector, pMK1001 or its derivative plasmid producing wild-type FliM, FliM_ΔC10_, FliM_ΔC20_, FliM_ΔC30_, and FliM_ΔC36_ was spotted onto VPG soft-agar plate and incubated at 30 °C for 6 h (A) and 8 h (B). (C and D) The rotational direction rates measured as in Fig. 4A. Cells prepared as (A) and (B) were observed. was spotted onto a TB soft-agar plate containing the final concentration of 0 mM, 1 mM or 10 mM IPTG, and incubated at 30 °C for 6 h.

### Effect of the expression of ZomB in *E. coli*

It is known that the overproduction of *S. putrefaciens* ZomB increases the switching frequency of the flagellar rotation of *E. coli* (36). We attempted to replicate a similar effect using ZomB from *V. alginolyticus*. We produced ZomB from a plasmid in *E. coli* wild-type strain (RP437), and performed the motility assay in the soft-agar plate (Fig. S3). We expressed *zomB* by the addition of 0, 1 and 10 mM IPTG in the plate, and found that the size of the motility ring was decreased in the 10 mM IPTG condition (Fig. S3), suggesting that motility was affected by the overproduction of ZomB from *V. alginolyticus*, similar to the case of ZomB from *S. putrefaciens*.

## Discussion

The rotational direction of the bacterial flagellar motor changes due to the conformational change of the C-ring. The conformational change is induced by the binging of CheY-P to the C-ring. ZomB is a single-TM protein specific for *Vibrio* spp. and *Shewanella* spp. and the Δ*zomB* mutant fixes the flagellar rotation in the CCW direction, similar to the Δ*cheY* mutant (36). The FliM-R49P mutant and FliG-G215A mutants confer CW rotation without CheY, indicating that C-ring structures can be fixed to the CW state without the CheY binding (42, 43). We found that ZomB was not necessary to maintain the CW state of FliM-R49P or FliG-G215A (Fig. 4 and 5). On the other hand, the cells with FliG-Q147H and FliG-A282T increased the CCW swimming by the Δ*zomB* mutation. As FliM-R49P and FliG-G215A mutations behave in CW swimming even in the *cheY* mutant and FliG-Q147H and A282T mutations behave in CCW swimming in the *cheY* mutant, we speculate that ZomB induces or maintains the CW state of the C-ring, which is made from the binding of CheY-P.

When a repellent signal such as phenol is given, the flagellar rotation of *Vibrio* spp. is not fixed to CW, but forward and backward movements are frequently repeated. This phenotype differs from those of *E. coli* and *Salmonella*. This suggests that the CW state structure of *Vibrio* spp. is very unstable, and quickly returns to the CCW state structure. In other words, the CheY-P binding is very weak and easily dissociates from the C-ring or the binding does not stably change the C-ring conformation in *Vibrio* spp. The binding property of CheY may be different between *E. coli*/*Salmonella* and *Vibrio* spp., and thus *Vibrio* spp. may need *Z*omB to support the CW state of the motor. The diameter of the C-ring is different between CCW and CW state (12, 13), the rotating pentamer of the A-subunits in the stator may engage in the C-ring like gears (11), and the rotational direction of the C-ring changes to either CCW or CW depending on the relative positions between the stator and the far side and the near side of the C-ring. If this model is correct, ZomB probably stabilizes the size of the C-ring in the CW state.

The CCW-locked phenotype by Δ*zomB* was suppressed by the deletion of the C-terminal 14 amino acid residue of FliM in *S. putrefaciens* (36). However, it was not suppressed in *V. alginolyticus*. This might reflect the difference in C-ring stability between *V. alginolyticus* and *S. putrefaciens* or that the conformational change caused by the *zomB* deletion differs between the two species. In addition, deletion of KKKW (251-254) motif, which is a highly conserved sequence at the C-terminus of ZomB, completely lacks the ZomB function in *S. putrefaciens*, but not in *V. alginolyticus*. This difference might also be explained by the difference in the stability of the C-ring between the two bacteria, and the KKKW motif might be necessary to maintain the ZomB structure in the membrane.

The candidate for the interaction partner(s) of ZomB might be FliG, FliM, CheY, PomA, or other unknown protein(s). FliG, FliM, and CheY are proteins whose mutants change or fix the direction of the flagellar rotation. As ZomB is predicted to be a one-TM protein, the protein should be present near the inner membrane. Our purification data suggest that ZomB forms multimeric complexes without any other proteins. It is possible that the ZomB multimer surrounds the C-ring like a housing of a motor to work as a spring to destabilize the diameter of C-ring to interact with FliG, FliM, PomA, or CheY (Fig. S4). At present, there is no evidence of what proteins are most likely to interact with ZomB. We will next investigate the protein that interacts with ZomB.

## Materials and Methods

### Bacterial strains, media and plasmids

The bacterial strains and plasmids used in this study are listed in Table S1. *V. alginolyticus* was cultured at 30 °C in a VC medium [0.5% (w/v) polypeptone, 0.5% (w/v) yeast extract, 0.4% (w/v) K_2_HPO_4_, 3% (w/v) NaCl, and 0.2% (w/v) glucose] or a VPG medium [1% (w/v) polypeptone, 0.4% (w/v) K_2_HPO_4_, 3% (w/v) NaCl, and 0.5% (w/v) glycerol]. When needed, chloramphenicol and/or kanamycin were added at final concentrations of 2.5 μg mL^−1^ and/or 100 μg mL^−1^, respectively, for *V. alginolyticus* culture. *E. coli* was cultured in an LB medium (Lennox; Nacalai Tesque). When needed, ampicillin, chloramphenicol or kanamycin was added at final concentrations of 50 μg mL^−1^, 25 μg mL^−1^ or 50 μg mL^−1^, respectively, for the *E. coli* culture.

*E. coli* was performed by a standard method using CaCl_2_. *V. alginolyticus* was transformed by the electroporation method described previously (44).

### Cloning of the *zomB* and *cheY* genes and preparing the *zomB* mutant

To construct pNT73, *zomB* DNA was amplified from the chromosomal DNA of VIO5 through PCR using a forward primer containing a BamHI site (CG<u>GGATCC</u>AGAAAAGTAAAGCAGACGCAAAGATAT) and a complement primer containing the PstI site (AA<u>CTGCAG</u>TTACCACTTTTTCTTCTCGCCGAAG) were cloned into the pMMB206 vector using the restriction enzymes and T4 DNA ligase (New England Biolabs). The *his_6_* gene was introduced using QuikChange site-directed mutagenesis (Agilent Technologies) using primers (G<u>CACCATCATCACCATCAC</u>AATATTGGTTTAATAATCGCTTTGG and <u>GTGATGGTGATGATGGTG</u>CATATCTTTGCGTCTGCTTTAC for *his_6_*-*zomB*, and G<u>CACCATCATCACCATCAC</u>TAACTGCAGCCAAGCTTGG and <u>GTGATGGTGATGATGGTG</u>CCACTTTTTCTTCTCGCCG for *zomB*-*his_6_*).

To construct pNT78, a *zomB* DNA fragment from the chromosomal DNA of VIO5 was amplified through PCR using a forward primer containing an NdeI site (GGAATTC<u>CATATG</u>CAGTATAAGGTCAAGGTAGAGA) and a complement primer containing the PstI site (CG<u>GGATCC</u>TTACCACTTTTTCTTCTCGCCGAAG) were cloned into the pET15b vector by using the restriction enzymes and T4 DNA ligase.

To construct pNT79, the upstream DNA (500 bp) and downstream DNA (478 bp) of the *zomB* gene were amplified through PCR using four primers containing the EcoRI site (G<u>GAATTC</u>CAGCAGGTGTCACCTGTGTTAAGTGC, AGACGCAAAGATCATTCACTTAAGGTCAATAAAGG, ACCTTAAGTGAATGATCTTTGCGTCTGCTTTACTT and G<u>GAATTC</u>GTGTAACTCCTCTTTAATCAAAAGCACC), were cloned into the pSW7848 vector using the restriction enzyme and T4 DNA ligase, as described previously (45).

To construct pHIDA6, the *cheY* DNA was amplified from the chromosomal DNA of VIO5 by PCR using a forward primer containing a BamHI site (AA<u>GGATCC</u>AGGTACCGGACACAAAATGACTAAAAACAC) and a complement primer containing EcoRI site (G<u>GAATTC</u>CCTGCAGTTATAAACGCTCGAAAATTTTATC) were cloned into the pTY60 vector by using restriction enzymes and T4 DNA ligase.

To delete the *zomB* genes in *V. alginolyticus*, pNT79 was introduced into *V. alginolyticus*, and homologous recombination was conducted as previously described (45).

### Motility assay in soft-agar plates

For *V. alginolyticus*, 2 μL of overnight cultures were spotted onto VPG soft-agar plates [VPG containing 0.25% (w/v) agar], and incubated at 30 °C for various time periods as described in the text. For *E. coli*, 2 μL of overnight cultures were spotted onto TB soft-agar plates [1% (w/v) Tryptone and 0.5% (w/v) NaCl containing 0.25% (w/v) agar], and incubated at 30 °C for various time periods. When needed, arabinose and isopropyl β-D-1-thiogalactopyranoside (IPTG) was added to the plate at final concentrations of 0.02% (w/v) and 1 or 10 mM, respectively, to induce the arabinose promoter and T7 promoter on the plasmid.

### Analysis of rotational direction and switching events

The rotational direction and switching events of the motor were observed as described previously (46) with slight modifications. Overnight cultures were diluted by 1/100 in VPG medium, and incubated with shaking at 30 °C for 4 h. When needed, arabinose was added into the VPG medium at a final concentration of 0.02% (w/v) to induce the arabinose promoter on the plasmid. After incubation, the cells were washed twice with V buffer [50 mM Tris-HCl (pH 7.5), 5 mM MgCl_2_, and 300 mM NaCl], and diluted by half in the buffer with or without 10 mM phenol, so that the final concentration of phenol was 5 or 0 mM, respectively. Cell motility was observed using high-intensity dark-field microscopy. We measured the rotational direction and switching events of the swimming cells for 10 s. The direction of the flagellar rotation was estimated from the position of the flagellum and the direction of the swimming; the flagella push the cell body during CCW rotation and pull it during CW rotation. The average values with standard deviations from more than 10 independent experiments are plotted in the graphs.

### Purification of ZomB_C_

The BL21(DE3) *E. coli* strain carrying pNT78 was cultured in LB broth containing ampicillin at 37 °C to an optical density at 660 nm of 0.6 to 0.8. IPTG was subsequently added to a final concentration of 0.5 mM and the culture was prolonged for about 20 h at 18 °C. Cells were collected by centrifugation (6,700 *g*) and suspended in TN buffer [50 mM Tris-HCl (pH 8.0), and 200 mM NaCl] containing cOmplete EDTA-free protease inhibitor (Roche) and lysozyme (Wako). The cells were then disrupted by sonication and centrifuged at 20,000 *g* for 10 min to remove cell debris. The supernatant was ultracentrifuged at 100,000 *g* for 30 min to remove the membrane and macromolecular complexes. The supernatant was loaded onto a column filled with Ni-NTA agarose (Qiagen), and the protein-bound agarose was washed with TN buffer containing 50 mM imidazole, and the proteins were subsequently eluted with TN buffer containing 400 mM imidazole. The eluted protein solution was mixed with thrombin to cleave the N-terminal His-tag, and dialyzed in TN buffer at 4 °C for approximately 14 h. The protein solution was loaded onto a column filled with Ni-NTA agarose and flow-through was collected to remove uncleaved proteins. The protein was concentrated using an Amicon Ultra 10 K device (Merck Millipore), loaded on a size exclusion column [ENrich SEC 650 10 x 300 Column (Bio-Rad)], and eluted with TN buffer using ÄKTA explorer (GE Healthcare). The ZomB_C_ protein-containing fractions were collected and concentrated using an Amicon Ultra 10 K device. The purity of the proteins was determined using SDS-PAGE.

### Crystallization and X-ray diffraction measurement

The purified proteins were crystallized using the sitting-drop vapor-diffusion method. The crystallization drops were prepared by mixing 0.5 μL of approximately 10 mg mL^−1^ protein solutions with 0.5 μL of the reservoir solutions. Initial screening was done using the following screening kits: Wizard Classic I and II, Wizard Cryo I and II (Rigaku), and Crystal Screen I and II (Hampton Research), and then the conditions were optimized. Crystals appeared within several weeks. The crystals used for the X-ray diffraction data collection were grown at 20 °C from drops prepared by mixing 0.5 μL protein solution (ca. 10 mg mL^−1^) in TN buffer with an equivalent volume of reservoir solution containing 30 % (w/v) PEG-4000, 100 mM Tris-HCl (pH8.5) and 200 mM MgCl_2_ for crystal 1, and 30 % (w/v) PEG-4000, 100 mM Tris-HCl (pH8.5) and 200 mM NaOAc for crystal 2.

X-ray diffraction data were collected at the synchrotron beamline BL41XU and BL45XU in SPring-8 (Sayo-cho, Japan). The crystals were frozen in liquid nitrogen and mounted in a nitrogen gas flow at 100 K for the X-ray diffraction data collection. The diffraction data were processed using *MOSFLM* (47) and were scaled with *AIMLESS* (48). The statistics of the diffraction data are summarized in Table S2.

## Acknowledgements

The synchrotron radiation experiments were conducted at the BL41XU and BL45XU of SPring-8 with the approval of the Japan Synchrotron Radiation Research Institute (JASRI) (No. 2018A2569, 2018B2569, 2019A2551, 2019B2551, and 2020A2574). This study was supported in part by JSPS KAKENHI Grant Numbers JP16J01859 (to N.T.), JP20J00329 (to T.N.), and JP20H03220 (to M.H.).

## Supporting information

Supplementary information associated with this article can be found in the online version available at the publisher’s website.

